# TWIST1 homodimers and heterodimers orchestrate lineage-specific differentiation

**DOI:** 10.1101/672824

**Authors:** Xiaochen Fan, Ashley J. Waardenberg, Madeleine Demuth, Pierre Osteil, Jane Sun, David A.F. Loebel, Mark Graham, Patrick P.L. Tam, Nicolas Fossat

## Abstract

The extensive array of bHLH transcription factors and their combinations as dimers underpin the diversity of molecular function required for cell type specification during embryogenesis. The bHLH factor TWIST1 plays pleiotropic roles during development. However, which combinations of TWIST1 dimers are involved and what impact each dimer imposes on the gene regulation network controlled by TWIST1 remain elusive. In this work, proteomic profiling of human-TWIST1 expressing cell lines and transcriptome analysis of mouse cranial mesenchyme have revealed that TWIST1 homodimer and heterodimers with TCF3, TCF4 and TCF12 E-proteins are the predominant dimer combinations. Dimers formation or their balance are altered by disease-causing mutations in TWIST1 helix domains, which may account for the defective differentiation of the craniofacial mesenchyme observed in patients. Functional analyses of the loss and gain of TWIST1-E-protein dimer activity have revealed previously unappreciated roles in guiding lineage differentiation of embryonic stem cells: TWIST1-E-protein heterodimers activate the differentiation of mesoderm and neural crest cells which is accompanied by epithelial-to-mesenchymal transition, while TWIST1 homodimers maintain the stem cells in a progenitor state and block entry to the endoderm lineage.

## Introduction

The helix-loop-helix (HLH) superfamily is an ancient group of transcription factors that appeared in unicellular organisms >600 million years ago and has continuously evolved and diversified since then to carry out ever-elaborating activities required for lineage specification (Murre, 2019). Through dimerization with different protein partners, bHLH factors drive the specification of many cell types during neurogenesis, haematopoiesis and myogenesis (Schlaeger *et al*., 2004; Belle and Zhuang, 2014; Comai and Tajbakhsh, 2014; Dennis *et al*., 2019). Eukaryotic HLH factors can be classified based on their structural and functional attributes (Massari and Murre, 2000). Class I, II and V factors are known to engage in homo- and heterodimerization. Class I proteins, comprised of all E-proteins (e.g. E12/TCF3, HEB/TCF12 and E2-2/TCF4), are broadly expressed in various tissues (Wang and Baker, 2015). Class II proteins are generally present in specific cell lineages, for example MYOD in myogenic cells, NEUROD in neurogenic cells and TWIST1 in mesenchymal tissues (e.g. craniofacial and limb mesenchyme; (Thisse *et al*., 1987; Lee *et al*., 1995; Spicer *et al*., 1996; Ghouzzi *et al*., 2000; Bildsoe *et al*., 2009; Bildsoe *et al*., 2013; Bildsoe *et al*., 2016). Class V represents a group of non-DNA binding HLH proteins, such as the ID proteins, that compete for E-protein binding (Benezra *et al*., 1990). It was hypothesised that the bHLH dimer composition is regulated by the expression level of each individual bHLH factor, their level of phosphorylation and their proportion relative to each other (Centonze *et al*., 2004; Firulli *et al*., 2005; Firulli *et al*., 2007). However, the diversity of dimer combinations and their functional specificity during development remains enigmatic.

The bHLH factor TWIST1 is highly expressed in cranial mesoderm- and neural crest-derived mesenchyme (Brault *et al*., 2001; Bildsoe *et al*., 2009; Bildsoe *et al*., 2013) where it is critical for craniofacial development. *Twist1*^*+/-*^ mice display craniosynostosis (Carver *et al*., 2002; Connerney *et al*., 2006) that partly phenocopies skeletal defects associated with *TWIST1* haploinsufficiency in human Saethre-Chotzen Syndrome (SCS, AHC [MIM: 101400]). Conditional ablation of *Twist1* in the cranial mesoderm (CM) or in the cranial neural crest (CNC) leads to malformations of the cranium, facial skeleton, brain, cranial nerves and muscles (Chen and Behringer, 1995; Soo *et al*., 2002; Ota *et al*., 2004). At the cellular level, *Twist1* is required for maintaining the mesenchymal cell morphology and their potency for osteo-, chondro-, and adipogenesis (Brault *et al*., 2001; Bildsoe *et al*., 2009; Miraoui *et al*., 2010; Bildsoe *et al*., 2013). Previous studies have highlighted the differential functions of TWIST1 dimers in the osteogenic differentiation of the cranial sutural mesenchyme (Connerney *et al*., 2006; Connerney *et al*., 2008), which is mediated by their targeted action on FGF signalling (Guenou *et al*., 2005; Rice, 2005; Miraoui *et al*., 2010). For example, the TWIST1-TCF3 heterodimer promotes mesenchymal stem cell proliferation, while TWIST1-homodimer activates *FGFR2, OCN* and *BSP* expression for ossification. Identifying TWIST1 dimerization partners and their transcriptional targets in mesenchyme will therefore allow for a better understanding of the mechanisms of development regulated by TWIST1 and bHLH factors dimers.

In this study, the diversity and expression of dimerization partners of TWIST1 were determined by mass-spectrometry analysis, following immunoprecipitation of human-TWIST1 (hTWIST1) from mesenchymal cells, and cross-compared with *Twist1* co-expression analysis in mouse embryonic head tissues. The bimolecular fluorescence complementation (BiFC) assay was used to elucidate the balance between hetero- and homo-dimerization and to assess the potential impact of pathological mutations. Finally, to dissect the specific functions of each TWIST1-dimer and their immediate downstream targets, we genetically engineered embryonic stem cells (ESCs), in which the expression of different TWIST1-E-protein dimers could be tightly controlled, and analysed their ability to differentiate and migrate. By delineating TWIST1 molecular interactions, our work has revealed previously unappreciated layers of control in lineage determination and cellular behaviour: TWIST1-E-proteins heterodimers promote mesoderm and neural crest differentiation through epithelial-mesenchymal transition (EMT) while TWIST1 homodimer maintains a progenitor-like state and blocks entry to the endoderm lineage. Together, using recent quantitative approaches and engineered cell models, this study has generated new insights into an ancient group of bHLH factors, the regulation of their dimerization activity and their role in fine-tuning lineage specification and differentiation.

## Results

### Identification of bHLH partners of TWIST1 in the embryonic head mesenchyme

In order to identify potential candidates dimerizing with TWIST1 protein, we first focussed on genes co-expressed with *Twist1* in vivo by investigating tissues of the mouse embryonic head. Microarray analysis of CNC and CM cells sorted from heads of embryonic day (E)9.5 embryos of *Wnt1-Cre:GFP* and *Mesp1-Cre:GFP* transgenic mice, respectively (Bildsoe *et al*., 2016; Fan *et al*., 2016), revealed that 58 out of 158 known bHLH factors (Skinner *et al*., 2010) were expressed in the head mesenchyme (Supplemental Table S1). Twelve bHLH factors were significantly enriched in CNC or CM (Fig. 1A) and 46 were expressed in both tissues (Fig. 1A and Supplemental Table S1). Based on their known roles in craniofacial development, seven candidates were selected for validation including SIM2, TCF4, EBF1, EBF3, TAL1, TWIST2 and TCF3 (isoform of E2A, a known TWIST1 partner as the positive control). HA-tagged proteins (including HA-tagged GFP as negative control) expression constructs were transfected into Madin-Darby Canine Kidney (MDCK) cells that stably over-express hTWIST1 (referred thereafter as MDCK/hTWIST1-OE cells) and have previously been used to investigate the role of TWIST1 to induce mesenchymal phenotypes (Xue and Hemmings, 2013; Bildsoe *et al*., 2016). These factors were co-immunoprecipitated with TWIST1. Reciprocally, TWIST1 was co-immunoprecipitated with EBF1, EBF3, TCF4 and, to a lesser extent, TWIST2, SIM1 and TAL1 (Fig. 1B), demonstrating that all these factors can complex with TWIST1 and are potential dimer partners. Furthermore, analysis by *in situ* hybridization revealed that genes encoding EBF1, EBF3 and TCF4 were co-expressed with *Twist1* in the frontonasal tissues, the cranial mesoderm and the first branchial arch of the mouse embryo (Fig. 1C).

**Fig. 1.**
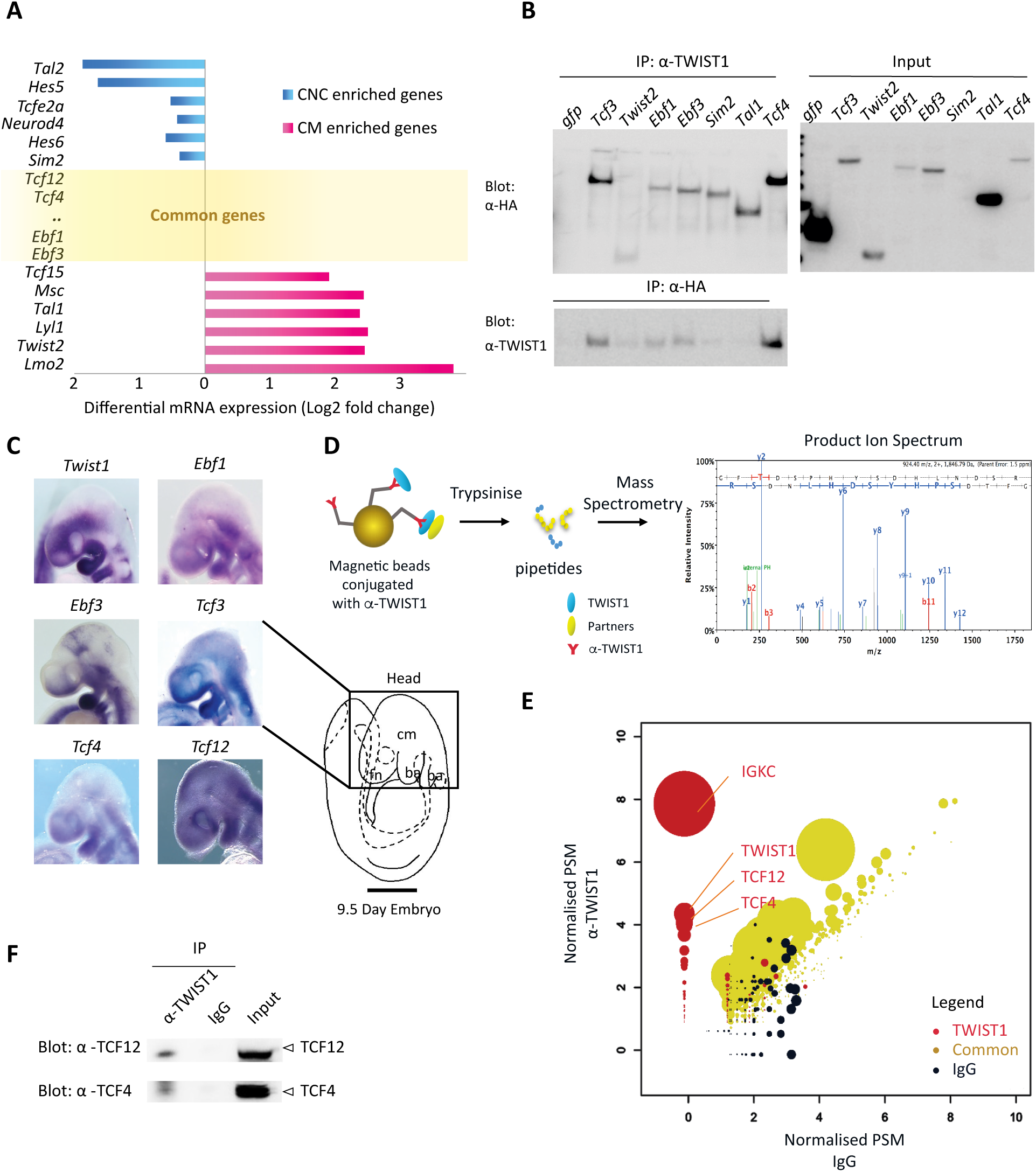
Unbiased high-throughput identification of proteins interacting with TWIST1 by co-expression analysis, co-immunoprecipitation (IP) and mass spectrometry. **A**. Relative expression of genes encoding known bHLH factors in cranial neural cells (CNC) versus cranial mesoderm (CM) of E9.5 embryonic head collated from transcriptome data previously published (Bildsoe *et al*., 2016; Fan *et al*., 2016). **B**. Detection of HA-tagged proteins (left blot: α-HA) after IP of TWIST1 (IP: α-TWIST1) and reciprocal IP of HA-tagged proteins (IP: α-HA) and detection of TWIST1 (Blot: α-TWIST1) from lysates of hTWIST1-expressing MDCK cells transfected with constructs expressing the HA-tagged protein partners, with expression confirmed in the input (right blot: α-HA) **C**. Expression of candidate transcripts analysed by whole mount *in situ* hybridization in regions of E9.5 embryonic head. Lateral views with anterior to the left. cm, cranial mesoderm; fn, frontonasal tissue; ba, brachial arch. Scale bar: 500 μm. **D**. Experimental strategy for identifying TWIST1 interacting partners from hTWIST1-expressing MDCK cells. TWIST1-containing protein complexes are isolated on beads using an α-TWIST1 antibody before analysed by tandem mass spectrometry which determines the ion spectrum of the different peptides (one example shown here). **E**. Groups of proteins identified using the non-parametric rank product method based on cumulative count of reporting a protein across independent experiments. Sets: red, TWIST1 IP; black, non-specific IgG; yellow, common to TWIST1 IP and IgG control. Normalised unique peptide spectrum match (PSM) of the detected proteins are shown on both axis for each condition. The size of the dots corresponds to –log_10_ P-value. IGKC: Immunoglobulin Kappa Constant **F**. Detection of TCF12 (Blot: α-TCF12) or TCF4 (Blot: α-TCF4) endogenous proteins after IP of TWIST1 (IP: α-TWIST1) from lysates of embryoid bodies over-expressing TWIST1, compared with results obtained for non-specific IgG and the input.

### Proteomic screening of MDCK/TWIST1-OE cells revealed TCF factors as the prevalent partners of TWIST1

Complementary to the above approach, TWIST1 dimer partners were identified through unbiased proteomic screening using affinity purification of the TWIST1-complexes from MDCK/hTWIST1-OE cells followed by mass spectrometry (Fig. 1D). Three biological replicates of immunoprecipitation using an α-TWIST1 antibody were performed in parallel with mouse IgG control. 846 proteins were identified and divided into three sets using the ranked-product method for differential analysis ((Breitling *et al*., 2004), Fig. 1E and supplemental Table S2). “Set 1” represents 103 identified proteins that specifically bound to TWIST1, “Set 2” are the proteins that only bound to IgG (377) and “Set 3” contains proteins bound to both IgG and TWIST1 (362). As expected, TWIST1 was one of the top enriched proteins in “Set 1” (Fig. 1E). Only two other bHLH factors, TCF12 and TCF4, were also highly enriched. The rest of “Set 1” TWIST1 putative partners included RNA-binding proteins and transcription factors such as Cyclin T2, TRIP6, GATA1, FHL2 and CRTC2 (Supplemental Table S2). *Tcf4* and *Tcf12*, like *Tcf3*, are expressed in the facial primordia and within *Twist1* expression domains in the frontonasal, the branchial arches and the mesoderm in the embryonic head (Fig. 1C). To recapitulate early development *in vivo*, we engineered mouse ESCs in which TWIST1 expression is inducible and differentiated these ESCs into embryoid bodies (EBs). Co-immunoprecipitation experiments performed with these EBs demonstrated that TWIST1 interacted with endogenous TCF4 and TCF12 (Fig. 1F).

### Mutations of conserved residues in bHLH domain impaired TWIST1-TCF12 dimer formation and function

The TWIST1 bHLH dimerisation domain shows 84–100% sequence identity among vertebrate orthologues (human, mouse, frog and zebrafish; Fig. 2A). It is also highly conserved (∼70% similarity) among several TWIST1 family proteins: TWIST1, TWIST2, HAND1 and HAND2, suggesting functional importance (Castanon and Baylies, 2002; Qin *et al*., 2012). There is a frequent incidence of clinically significant point mutations (63 reported mostly in SCS patients) in the bHLH domain of *TWIST1* ((Fig. 2A; Human genomic mutation database; (Stenson *et al*., 2003; Centonze *et al*., 2004; Firulli *et al*., 2005; Firulli *et al*., 2007; Barnes and Firulli, 2009). Some of these mutations are located at or around the threonine and serine phosphosites (Fig. 2A) and could inhibit TWIST1 phosphorylation (Firulli *et al*., 2005; Firulli and Conway, 2008). It was previously shown that TWIST1 SCS mutations in the basic-helix I region, such as R118H, may impair phosphorylation at T121 and S123 residues and alter the preference of dimerization with TCF3 and HAND2 (Firulli *et al*., 2005).

**Fig. 2.**
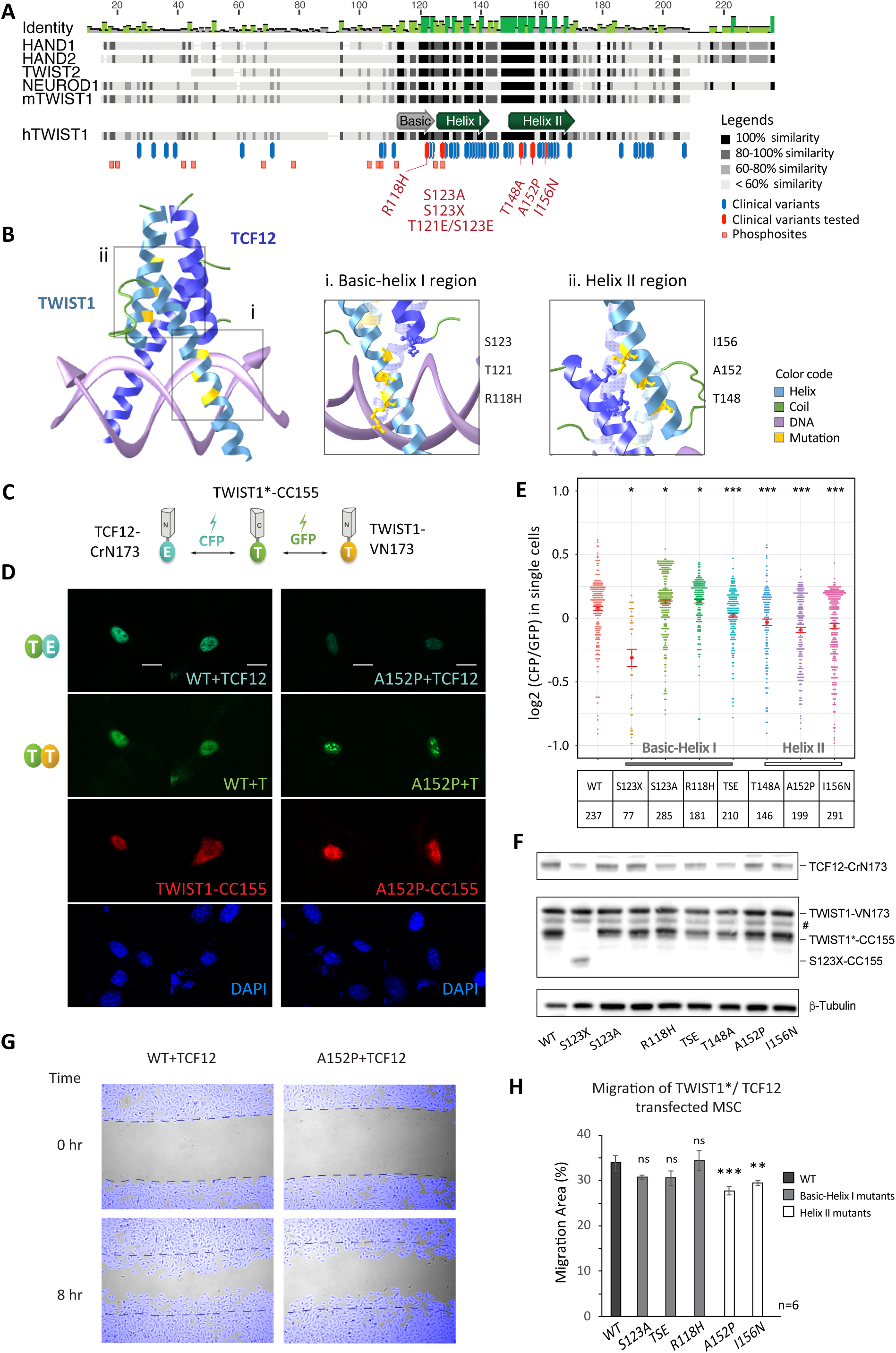
Human disease-causing mutations disrupt TWIST1-TCF12 dimer function. **A**. Alignment of selected mouse TWIST-family bHLH proteins and hTWIST1 sequences. The degree of similarity of the different regions is indicated in dark shades. Blue and red bars indicate missense pathological mutations reported in the Human Gene Mutation Database (Stenson *et al*., 2003). The red bars identify the mutations analysed in this study. The pink squares are predicted phosphosites. **B**. *In silico* 3D structure prediction of the bHLH regions of TWIST1-TCF12 dimer binding to DNA, based on the structure of NeuroD-TCF3 dimer as determined by crystallography (O’Donoghue et al. 2015; http://aquaria.ws). Enlarged representation of **i**. basic-helix I and **ii**. helix II domains of TWIST1 are shown. TWIST1 residues mutated in disease and their atomic interaction with TCF12 are highlighted in yellow. **C**. Experimental setup for the BiFC assay for which three independent constructs expressing TWIST1*-CC155, TWIST1-VN173 and TCF12-CrN173 were equimolarly co-transfected in C3H10T1/2 cells. TWIST1*: wildtype or SCS variants. Interactions between the different proteins and the relative proportion of TWIST1 heterodimers versus TWIST1 homodimers are then assessed by quantifying and comparing CFP and GFP fluorescence intensities as shown in E. **D**. Representative confocal microscopy images of BiFC assays performed for wildtype (WT) and A152P-mutant TWIST1. Signals for dimers: TWIST1*-CC155/TWIST1-VN173 (T/T) or TWIST1-VN173/TCF12-CrN173 (T/E) were shown as indicated. Red channel shows TWIST1*-CC155 detected by immunostaining with an α-HA antibody. Cell nuclear was stained by DAPI. Scale bar: 10 μm. **E**. Heterodimer/homodimer CFP/GFP ratio quantified for individual cells (each dot corresponding to one cell) in BiFC assays. TWIST1 variant and number of cells assayed are indicated in chart below. TSE corresponds to T121E; S123E double mutant. Red bars represent standard errors of the mean. Data analysed by non-parametric Mann-Whitney U test (two-tailed); *P ≤ 0.05, *** P ≤ 0.001. F. Western blot analysis of BiFC proteins expression in C3H10T1/2 cells. TWIST1 BiFC constructs were detected with an α-GFP antibody; TCF12-CRN173 was detected with an α-TCF12 antibody. Loading control (β-tubulin) is also shown. #: non-specific band. **G**. Example images from a scratch assay. C3H10T1/2 cells were co-transfected with TWIST1*-CC155 (WT or A152P) and TCF12-CrN173 expression vectors. Time lapse images were taken every 15 mins over an eight-hour-period, after a scratch was made on the cell monolayer. Cells were tracked (pseudo-coloured in blue) and area of cell free gap was quantified in ImageJ software. Dashed lines indicate original gap boundaries. **H**. Quantification of migration (% of total area in 8 hrs), of C3H10T1/2 cells co-transfected with TWIST1*-CC155 and TCF12-CrN173 expression vectors. Data represent the mean ± standard errors of n=6 independent experiments, analysed by one-way ANOVA with Holm-Sidak’s post-test; **P ≤ 0.01, *** P ≤ 0.001 or non-significant (ns).

To validate the putative interactions identified above and test the hypothesis that TWIST1 pathological mutations in the bHLH region may shift the balance of homo-versus hetero-dimerizations, we employed the BiFC assay, a bimolecular complementation system which allows the detection of direct physical interaction between two partners by fluorescence (Hu and Kerppola, 2003). A subset of mutations with known impact on the basic-helix I region was selected (Fig. 2A, red): truncation mutation in the bHLH domain (S123X), phosphorylation-inhibitory mutations (S123A and R118H) and phospho-mimetic mutations (T121E and S123E). We also selected previously uncharacterised mutations in helix II (T148A, A152P and I156N) that affect conserved residues among TWIST-family members. When examined by 3D *in silico* modelling of the bHLH regions of a TWIST1/TCF12 dimer in the presence of DNA, the phospho-regulatory residues appear to be in close proximity to the DNA interface rather than the site in contact with the E-protein (Fig. 2Bi) while clinical variants on helix II affect residues located at the protein interaction interface (Fig. 2Bii).

We tested the impact of these *TWIST1* mutations on TWIST1 ability to dimerize with TCF12, which is also associated with SCS and other craniofacial anomalies (Sharma *et al*., 2013). TWIST1 and TCF12 were conjugated with different split CFP or GFP fluorescent domains (N- or C-terminal: CrN173 and VN173, and CC155, respectively) and co-expressed in C3H10T1/2 cells where dimerization was monitored by fluorescence (Fig. 2C). The interaction of TWIST1-CC155 with TCF12-CrN173 versus TWIST1-VN173 was measured by the ratio of fluorescent intensity from the two different fluorophores (Fig. 2D, E). When expressed at comparable levels (Fig. 2F), TWIST1 mutants displayed significant changes in their preference to form homo- (GFP) or heterodimers (CFP; Fig. 2D, E). Among all mutations tested, truncation at the bHLH domain (S123X) severely disrupted dimerization, while loss of the N-terminus did not affect dimerization (Fig. 2E and supplemental Fig. S1A). Phosphorylation-inhibitory mutations S123A and R118H increased the preference for heterodimer formation, whilst the phospho-mimetic mutant (TSE for T121E & S123E) of TWIST1 favoured homo-dimerization (Fig. 2E). Finally, all three missense mutations at helix II compromised more severely heterodimerization (Fig. 2E).

Mutations associated with significant shift in dimer balance were further tested for their potential impact on cell migration, a known TWIST1-driven biological process. Cell migration was tracked and quantified after a scratch was made on a confluent monolayer of mesenchymal stem cells transiently expressing TCF12 and different variants of TWIST1 (Fig. 2G). Mutations in helix I showed no effect on the cells, but cells expressing TWIST1 with helix II mutations (A152P, I156N) showed reduced cell migration (Fig. 2G-H). Altogether, these results demonstrate that residues in helix I and helix II are important determinants for dimerization, and that disrupting TWIST1/TCF12 dimerization with mutations at the interaction site in helix II may compromise mesenchymal migration.

### Loss of *Twist1* impairs exit from pluripotency and cell fate specification

During ESCs differentiation, the expression of endogenous *Twist1* was first detected from Day 3 and then peaked at Day 6-7 (Fig. 3A). We first generated and studied *Twist1* loss-of-function (LOF) ESCs to appreciate the overall function of the ensemble of TWIST1 dimers during differentiation. Mono- and bi-allelic *Twist1* knockout mouse ESCs were engineered using the CRISPR-Cas9 methodology (Fig. 3B, C), and the expression of pluripotency and lineage gene markers was analysed (Fig. 3D-H). Pairwise statistical tests revealed that both full and partial loss of *Twist1* disrupted lineage specification in a gene dosage-dependent manner. Genes associated with pluripotency such as *Dppa2, Nanog, Sox2* and *Pou5f1* were reduced in wildtype parental cell line at Day 3, but expression was retained at the pre-differentiated levels in the mutant cells (Fig. 3D). Concurrently, ectodermal and mesodermal genes failed to be activated in the mutant cells (Figure 3E, F). Expression of EMT genes such as *Pdgfra, Snail2* and *Vegfa* were also reduced in the mutant cells while the epithelial marker, *Cdh1*, was up-regulated (Fig. 3F). In contrast, endodermal markers such as *Gsc, Rbm47, Foxa2* and WNT signalling components (ligand: *Wnt3*, receptors: *Fzd10* and *Prickle1*, target gene: *Axin2*) were up-regulated in the mutant cells (Fig. 3G, H). These functions of *Twist1* in selectively directing lineage differentiation and maintaining the mesenchymal progenitors are likely to be orchestrated by a combination of TWIST1 dimers.

**Fig. 3.**
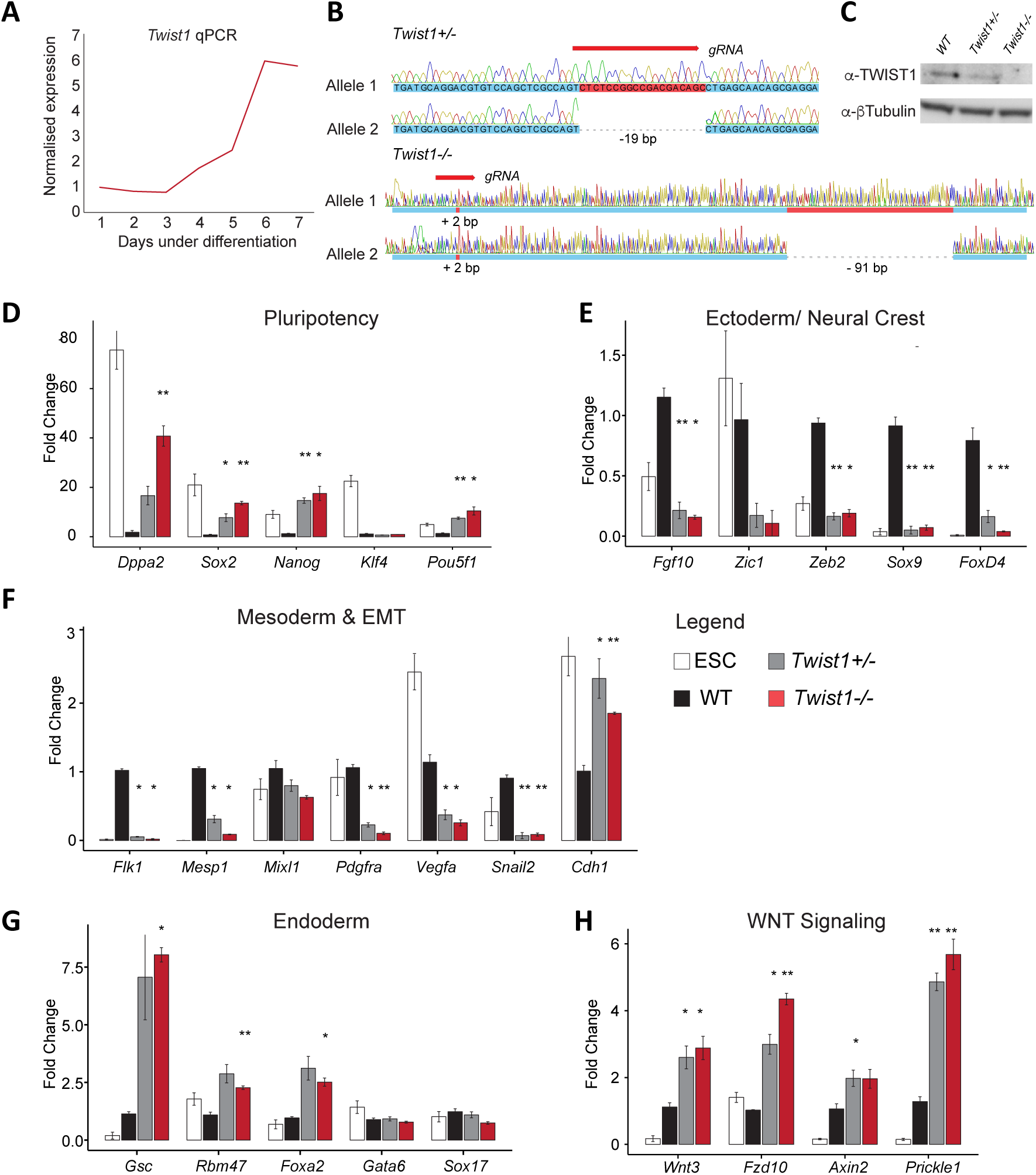
TWIST1 is required for mesoderm and ectoderm differentiation. **A**. RT-qPCR analysis of *Twist1* expression relative to mean value of *Gapdh* expressions (Y-axis) during ESC differentiation (X-axis). Each time point represents the mean value of n=3 independent experiments. **B**. *Twist1* alleles of two independent ESC clones (*Twist1*^*+/-*^ and T*wist1*^*-/-*^) generated by CRISP-Cas9 editing. Genomic positions targeted by the gRNA, Sanger sequencing chromatograms and deduced gDNA sequences are shown. Mutations are frameshifts in the coding sequence of exon 1. *Twist1*^*+/-*^ clone: allele #1 is unmodified; allele #2: 19 bp deletion. *Twist1*^*-/-*^ clone: allele #1: 2 bp insertion; allele #2: 2 bp insertion and 91 bp deletion. **C**. Western blot analysis of TWIST1 proteins in the clones characterised in B and the parental ES wildtype (WT) cell line. Loading control (β-tubulin) is shown. **D-H**. RT-qPCR analysis of the expression (Y-axis) of selected genes (X-axis) involved in cell lineage, EMT or WNT signalling in mutants, wildtype (WT) and undifferentiated ESC, at day 3 of *in vitro* differentiation. Expressions were normalized against the mean value of *Tbp* and *Gapdh* expressions and the wildtype sample at Day 3 of differentiation. Each data points represent the mean ± standard errors of n=3 independent experiments. Data analysed by non-parametric Mann-Whitney U test (two-tailed). *P ≤ 0.05, **P ≤ 0.01, ***P ≤ 0.001.

### Comparison of the target specificity that differs between different TWIST1 dimers

To deconstruct the function of each TWIST1 homodimer and heterodimers during lineage differentiation, a gain of function (GOF) approach was taken to analyse the downstream genes controlled by individual dimers. In *Drosophila*, tethered Twist-Twist and Twist-Dorsal dimers could recapitulate the function of dimers formed naturally by Twist1 and Dorsal. Twist homodimer induced *Mef2* expression and muscle differentiation *in vivo*, whereas the Twist-Dorsal dimer antagonised this activity (Castanon and Baylies, 2002). Following the same strategy, we tethered TWIST1 and TCF proteins into dimers via flexible poly-glycine linkers: TWIST1-TWIST1 (TT), TWIST1-TCF3 (T3), TWIST1-TCF4 (T4) and TWIST1-TCF12 (T12), and expressed them in ESCs (Fig. 4A-C). The expression constructs were stably integrated into a designated genomic locus of the A2LoxCre ESCs, downstream of a doxycycline inducible promoter (Mazzoni *et al*., 2011; Iacovino *et al*., 2014; Sibbritt *et al*., 2019). ESCs were differentiated for three days into EBs (Fig. 4B) and assayed for the expression of developmental/lineage marker genes. Before doxycycline induction, no expression of the transgene was detected (Supplemental Fig. S2A). Dimer expression was induced at Day 2 of differentiation in synchrony with the onset of expression of endogenous *Twist1* (see Fig. 3A). To ensure that the effects of each dimer was not confounded by dissimilar level of expression, different doses of doxycycline were tested and the optimal concentration was selected for each cell line to achieve comparable levels of expression between each construct (Fig. 4C and supplemental Fig. S2A).

**Fig. 4.**
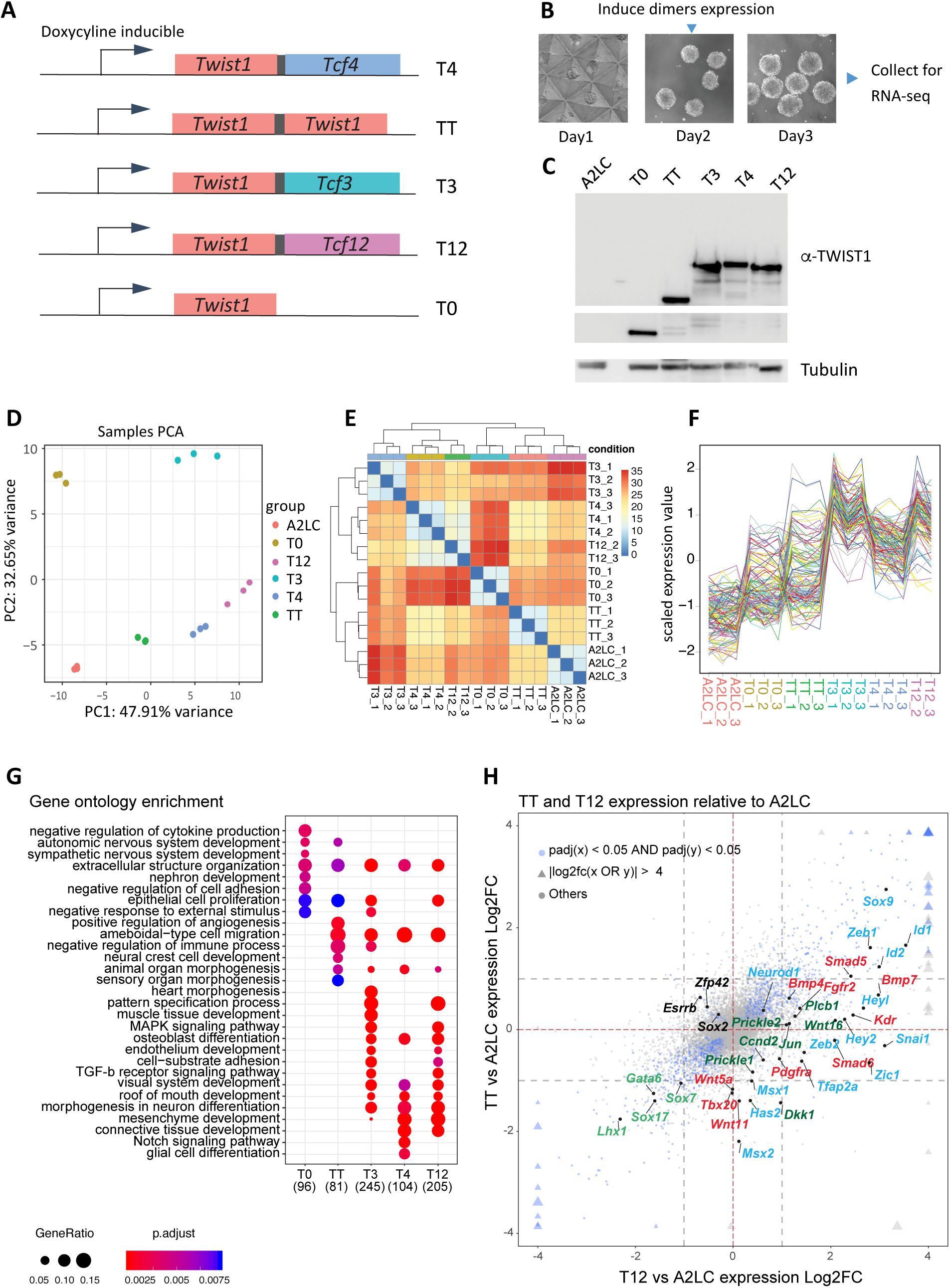
Transcriptome profiles of embryoid bodies expressing different TWIST1-dimers and monomer. **A**. Schematic of doxycycline-inducible expression constructs: *Twist1-Tcf3* (T3), *Twist1-Tcf4* (T4), *Twist1-Tcf12* (T12), *Twist1-Twist1* (TT) and *Twist1* only (T0; TWIST1 monomer) recombined into designated locus in A2loxCre parental ESCs (Mazzoni *et al*., 2011; Iacovino *et al*., 2014). Proteins in dimer are joined by a flexible poly-glycine linker (black box). **B**. Experimental setup to obtain dimers expressing EBs: ESCs are differentiated into EBs for 3 days. Dimer expression is induced with doxycycline 16 h before collection **C**. Detection by western blot using an α-TWIST1 antibody of TWIST1 dimers and monomer in the engineered ESCs and the parental A2loxCre cell line (A2LC) after doxycycline induction. Loading control (β-tubulin) is also shown. **D**. Principal component analysis of the transcriptomes of the dimers/monomer-expressing ESCs and A2LC EBs (n=3 independent RNA-Seq for each cell line) showing the first two primary components. **E**. Correlation map of the different RNA-seq data showing Euclidean distance between cells lines (from high correlation in blue to low correlation in red). **F**. Scaled expression (Y-axis) of the 125 genes with top PC loadings across all cell lines. Each line corresponds to one gene. **G**. Gene Ontology analysis of upregulated genes (adjp < 0.05, Log2FC > 1) in all cell lines versus A2LC. The Top 10 most significant non-redundant GO categories for each cell line were plotted. Dot sizes represents “GeneRatio” (gene upregulated/gene in GO set) and adjusted P-value is colour-coded. Number of identified genes in each group is show between parentheses. **H**. Four-way plot visualization of DESeq2 result. Log2FC comparisons between TT and T12 treatments versus A2LC were performed. Blue nodes indicate genes significantly changed for both cell lines. Triangular shapes indicate values exceeding the limit of for the plot. Size change reflects the magnitude of Log2FC values. Vertical and horizontal dashed lines indicate |Log2FC|= 1. Key markers and pathway genes are indicated: Black, pluripotency; Green, endoderm; Red, mesoderm; Blue, ectoderm & EMT; Dark green, WNT signalling pathway.

The transcriptomes of EBs differentiated for 3 days expressing comparable levels of dimers were then analysed by RNA-Seq and compared to the transcriptome of the parental A2loxcre cell line (A2LC) as well as the transcriptome of a cell line expressing TWIST1 Monomer (T0) that were both also differentiated for 3 days (Fig. 4D-H and supplemental Fig. S3). Unsupervised principle component analysis (PCA) and hierarchical clustering confirmed groupings of replicates (Fig. 4D, E). Global gene-expression profile separated T3, T4 and T12 from TT, T0 and A2LC EBs (Fig.4E, F), a result which was mainly driven by genes upregulated in the heterodimers. Among the three heterodimers, cells expressing T4 and T12 dimer exhibited more similar gene expression profile (Fig. 4E). Differentially-expressed-genes (DEGs) analysis was performed for T0, TT, T3, T4 and T12 against the A2LC cells. Examination of the upregulated gene groups revealed that while all dimers activated cell migration program, the heterodimers specifically activated the expression of genes associated with pattern specification, mesenchymal tissues development (muscle, skeletal, connective tissue etc.) and TGFβ signalling (Fig. 4G). A four-way comparison between TT or heterodimers against A2LC also demonstrated a stronger ability of the heterodimers to activate developmental markers, especially mesenchyme-related genes (Figure 4H; Supplemental Fig. S3).

It was previously demonstrated by *in vitro* mobility shift assay that different dimers preferentially bind different DNA sequences (Firulli *et al*., 2007; Chang *et al*., 2015). Sequence motif analysis was performed on the proximal regulatory sequences for dimer-specific DEGs. For each group, 100-600 genes were up-regulated, and 100-200 were down-regulated (adjusted p-value ≤ 0.05, fold change ≥ 2; Fig. 5A; Supplemental Table S3). Top ranking regulatory motifs for heterodimers-activated promoters contained a single canonical E-box “CATCTG” or “CAGCTG” (Fig. 5B), which is consistent with TWIST1 ChIP-seq and dimers mobility shift assays results (Firulli *et al*., 2007; Chang *et al*., 2015). TT targets were uniquely enriched in double E-boxes with a five nucleotide spacer as previously reported by ChIP-Seq in human mammary epithelial cells over-expressing TWIST1 (Chang *et al*., 2015).

**Fig. 5.**
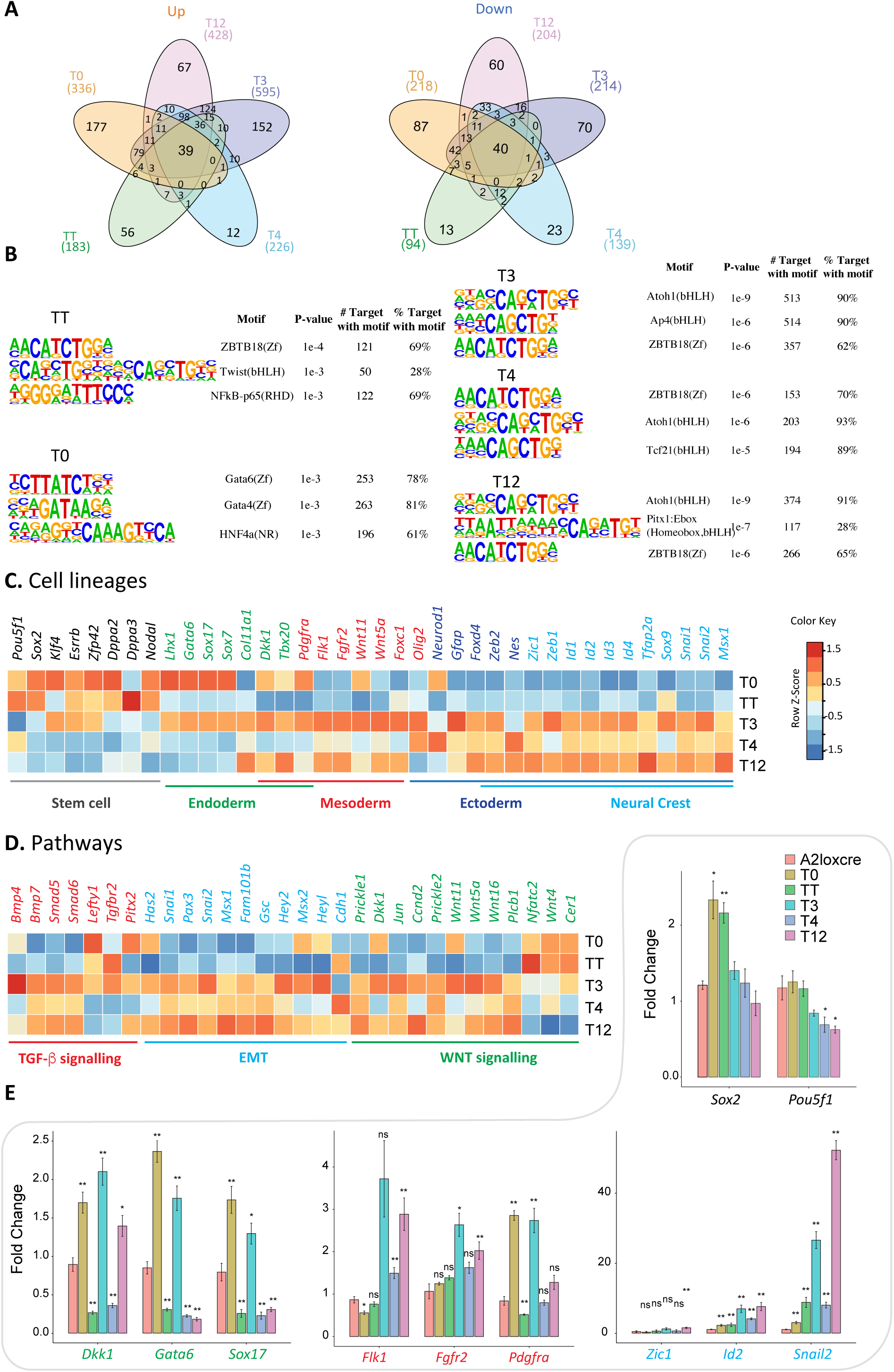
Differential specificity analysis of TWIST1-dimers and monomer. **A**. Pairwise differential gene expression analysis of the dimers/monomer-expressing ESCs against the parental A2LoxCre ESCs performed with DESeq2 (Love *et al*., 2014) (adjp < 0.05, fold change ≥ 2). Number of individual and common up-regulated (Up) and down-regulated (Down) genes are shown in Venn diagrams. **B**. Top three predicted binding motifs enriched in the regulatory sequences (TSS +/- 2 kb) of the differentially regulated target genes of TWIST1 dimers, generated with HOMER (Heinz *et al*., 2010). Names of the known motif and percentage of targets containing these motifs are listed, **D**. Heatmap showing the relative expression (Log_2_-fold vs A2LC) of selected genes involved in cell lineage, EMT or cell signalling pathways for the dimers/monomer-expressing EBs. Colour-coded normalised Z-score is shown. **E**. RT-qPCR analysis of the expression (Y-axis) of key lineage markers (X-axis) in different EBs. Expressions were normalized against the value of *Tbp* and the wildtype sample at Day 3 of differentiation. Each data points represent the mean ± standard errors of n=3 independent experiments. Data analysed by non-parametric Mann-Whitney U test (two-tailed). *P ≤ 0.05, **P ≤ 0.01 or non-significant (ns). Black, pluripotency markers; Green, endoderm markers; Red, mesoderm markers; Blue, neural crest markers.

To further dissect the functionality of each dimer, we performed additional enrichment analyses focusing on development related gene sets and taking into account gene expression changes relative to the parental A2LC cells. Neural-ectoderm differentiation derivatives were repressed by the monomer while mesoderm and endoderm cell fates were repressed by TT (Supplemental Fig. S4A; Supplemental Fig. S3; Supplemental Table S3). On the other hand, heterodimers activated overlapping sets of genes associated with mesoderm and neural ectoderm development (Supplemental Fig. S4A; Supplemental Table S3). T3 dimer activated heart developmental processes, including heart vasculature, muscle, and blood cells, while the most frequent processes induced by T4 were related to neuron differentiation (Supplemental Fig. S4A; Supplemental Table S3). T12 activity was intermediate between T3 and T4. A comparison down to key cell lineage and pathway genes also reflected these changes (Fig. 5C, D) which were furthermore validated by RT-qPCR experiments (Figure 5E). The most evident changes in the heterodimer groups were the downregulation of pluripotency genes and a burst of expression of lineage markers, especially those for mesoderm and neural crest ectoderm development (Fig. 5C, E). Endoderm markers were repressed by all dimers except T3 (Fig. 5D, E; Supplemental Fig. S3; Supplemental Table S3).

Taken together, our results have demonstrated that TWIST1 dimerization partners drive different gene expression programs which may be influenced by the preference for specific DNA motifs. Heterodimers appeared to drive ectoderm and mesoderm differentiation, whereas TWIST1 homodimer repressed the expression of developmental genes while maintaining the pluripotency state.

### TCF12-TWIST1 dimer activates EMT more robustly and antagonises TWIST1 homodimer activity for critical signalling pathway genes

*Twist1* and *Tcf12* have been shown to act synergistically for cranial tissue development (Sharma *et al*., 2013). Previous studies have suggested that homo- and heterodimers regulate antagonistically the skull suture (Connerney *et al*., 2006; Connerney *et al*., 2008). Therefore, we focused further our analysis on the T12 dimer and its ability to induce transcription of developmental pathways, in comparison with TT (Supplemental Fig. S4B, C and supplemental Table S4). T12 expressing cells significantly activated more genes associated with TGFβ signalling (FDR 0.007), WNT signalling (FDR 0.017) and EMT (FDR 0.002; around 50% of all EMT genes were enriched in T12 cells; Fig. 6B). T12 expressing cells were also enriched for genes associated with processes of neuronal, skeletal, face and cardiovascular system development (Fig. 6B and supplemental Fig. S4B). Examining specifically genes encoding transcriptional regulators associated with craniofacial development (Fig. 6B, Face development), we noted that a subset of critical developmental regulators such as those associated with TGFβ signalling (e.g. *Tgfbr1, Pdgfrα*) and WNT signalling (*Prickle1, Dkk1*) were antagonistically regulated by the homodimer and the heterodimer (Fig. 6C). Other genes, represented by *Fgfr2, Jun, Chd7* and *Mmp2* were activated to a greater extent by T12 than TT (Fig. 6C).

**Fig. 6.**
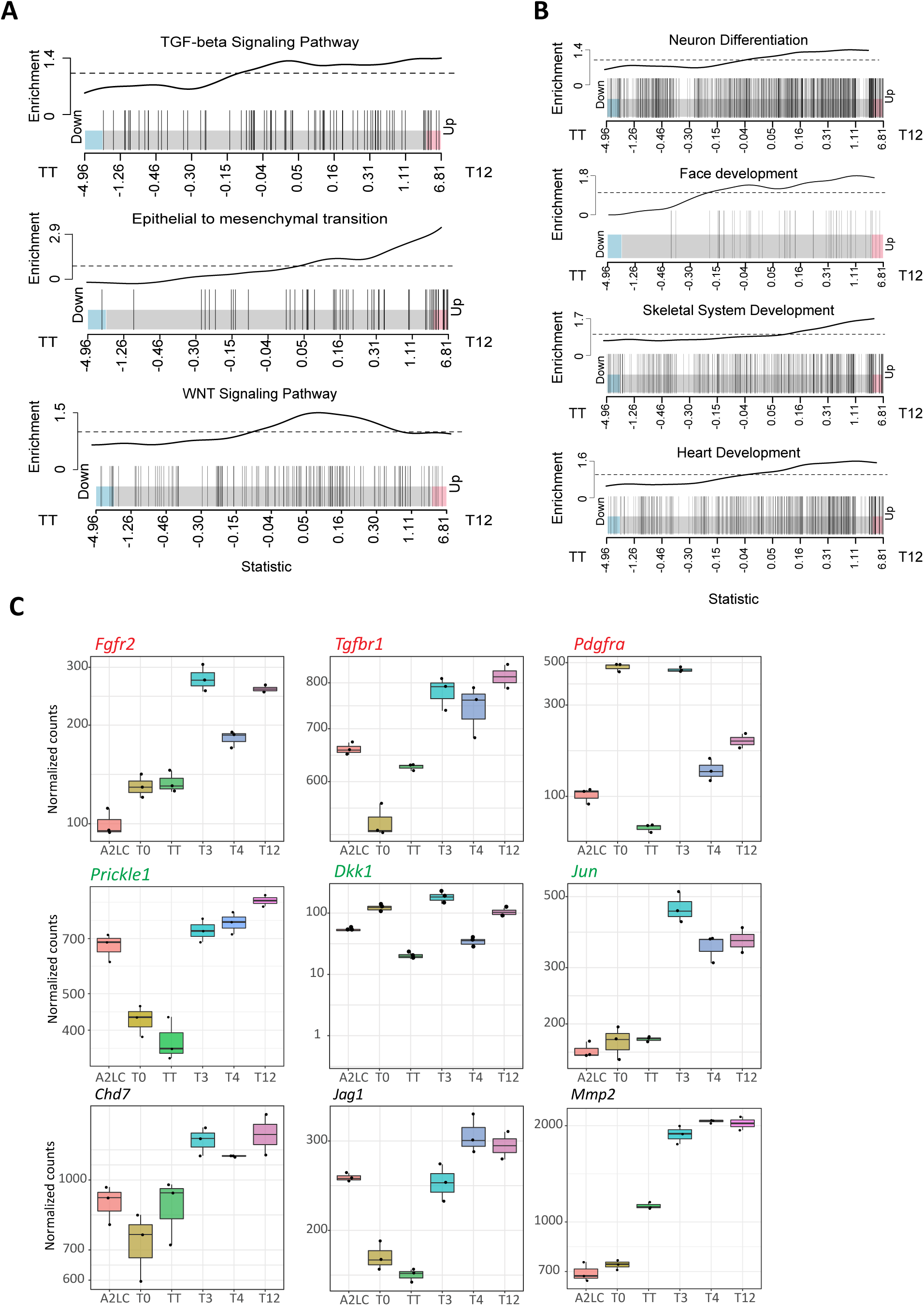
TWIST1-TCF12 heterodimer antagonises TWIST1 homodimer. **A, B**. Barcode plots of the enrichment of cell signalling and Developmental process-related genes in TWIST1-TCF12 EBs (right) versus TWIST1-TWIST1 EBs (left). Individual genes, represented by vertical bars, are ranked from left to right based on log_2_ fold change (X-axis) for TWIST1-TCF12 EBs versus TWIST1-TWIST1 EBs. The enrichment line shows the relative local enrichment (Y-axis) of the bars in the plot. The dotted horizontal line indicates neutral enrichment. Values above the dotted line show enrichment while values below the dotted line show depletion. **C**. Boxplots depicting the expression (Y-axis) of selected genes important for craniofacial development and cell signalling pathways that are enriched in TWIST1-TCF12 versus TWIST1-TWIST1 expressing EBs. In Red, TGFβ signalling related genes; In Green, WNT signalling related genes. Dots represents replicate data points. The top and bottom edge of each box shows first and third quantiles (Q1 and Q3) of the data values; the middle bar shows median values. Upper whisker = highest observed value within [Q3, 1.5*Q3]; Lower whisker = lowest observed value within [0.5*Q1, Q1].

The differential expression of downstream genes between T12 and TT and the functional consequences for craniofacial development are therefore likely to be the outcome of the balancing activity of these two dimers for transcriptional regulation.

## Discussion

### TCF factors are the major dimerization partner and transcriptional modulator of TWIST1

Our study has identified TCF3, TCF4 and TCF12 E-proteins, which are expressed in the cranial mesenchyme, as the predominant interacting partners of TWIST1. This is also supported by the results of a TWIST1 proximity labelling screen showing that the TCF factors partner with TWIST1 consistently in neural crest cells, mesenchymal stem cells and 3T3 cells (Fan et al, unpublished results). E-proteins are also major dimerization partners of several other Class II bHLH factors (Murakami *et al*., 2004; Forrest *et al*., 2014). For example, TCF3, TCF4 and TCF12 are top interacting candidates of HAND2 and enhance its transcriptional activity at the promoter of target genes (Murakami *et al*., 2004). Consistent with these observations, we found that TWIST1 induced differentiation programs more effectively when coupled with E-proteins. Also similar with HAND2, coupling TWIST1 with different E-proteins resulted in slightly altered DNA binding activity as previously shown by electromobility shift assay (Murakami *et al*., 2004; Firulli *et al*., 2007). These observations suggest that E-proteins might be the primary transcriptional modulators of TWIST1, and potentially other class II bHLH factors, for the regulation of various tissue-specific programs during development.

### Pathological mutations induce bHLH factor dimerization imbalance

For TWIST-family bHLH factors, dimerization can be modulated by phosphorylation of the bHLH domain (Centonze *et al*., 2004; Firulli *et al*., 2005; Firulli *et al*., 2007). Hypo-phosphorylation mutations (T121A, S123A) near or at the helix I phosphorylation sites were previously shown to favour homodimerization (Firulli *et al*., 2005). Nevertheless, a triple phospho-mutant model of TWIST1 (S123A, T148A and A184A) in oncogenic cell lines showed preference for dimerization with TCF3 and HAND2 (Wang *et al*., 2017). In line with this, we found that two individual hypo-phosphorylation mutants (R118H, S123A) both increased preference for TCF12 heterodimer formation, whilst phospho-mimetic forms at T121 and S123 reduced heterodimerization. We noted that incorporation of the negatively charged phosphate group to these residues may reduce binding to DNA, rather than directly interfering with the dimerization interface. Therefore, T121/S123 dephosphorylated TWIST1 may favour heterodimerization through stabilising DNA-binding or adapting a helix conformation that enhances E-protein binding. Driven by a potentially different mechanism, TWIST1 helix II mutations at the interaction site severely compromised hetero-versus homodimerization potential, which had greater functional consequence than helix I variants and impacted on the migration of mesenchymal cells. Altogether, these observations demonstrate that these residues constitute an important interface for dimer balance adjustment and therefore play a critical role for cell differentiation and development.

RNA-seq analysis of differentiating ESCs revealed that the TWIST1-TCF12 heterodimer is a potent driver of EMT and mesenchymal differentiation compared to TWIST1 homodimer. TWIST1-TCF12 also induced signalling pathways and key transcription regulators associated with head mesenchyme development. These genes were either inactive or repressed in cells expressing the TWIST1 homodimer. The *Twist1* heterozygous (*Twist1*^*+/-*^) mouse model that recapitulates SCS phenotypes displays excessive FGF and TGFβ signalling at the osteogenic front, as revealed by the expansion of FGFR2, pSmad 1/5/8, and ID protein signal (Connerney *et al*., 2006; Connerney *et al*., 2008). These mis-regulated signalling activities could therefore be a consequence of imbalanced TWIST1-E-protein activities.

### Dimerization-dependent activities of TWIST1 in progenitor cell lineage specification

TWIST1 regulates multiple pathways during lineage specification. Loss of *Twist1* function leads to retention of pluripotency, defective mesoderm and ectoderm differentiation, as well as disrupted EMT. EMT is tightly associated with ESCs differentiation, especially the specification of mesoderm (Nakaya and Sheng, 2008; Evseenko *et al*., 2010; Iacovino *et al*., 2014). Major EMT drivers including *Twist1, Zeb2, Snail1* and *Snail2* are induced by canonical WNT signalling in mesendoderm differentiation (Ten Berge *et al*., 2008), while constitutive activation of WNT further directs precursor cells into primitive endoderm (Price *et al*., 2013). Both *Twist1* and *Snail1* are required for WNT induced mesoderm differentiation (This study and (Sharma *et al*., 2013) but only *Twist1* negatively feedbacks on WNT signalling regulation and represses endoderm cell fates. Over-expression of the TWIST1 homodimer blocked WNT activity and endoderm differentiation while loss of TWIST1 shifted differentiation towards endoderm. On the other hand, ablation of *Snail1* promotes ectodermal instead of mesendodermal fate without affecting WNT pathway genes (Sharma *et al*., 2013). These observations suggest a dichotomic control carried out by TWIST1 during development: 1) an EMT-dependent role in promoting mesoderm and neural crest commitment and 2) an EMT-independent role in endodermal repression potentially by negatively feedback on WNT activity.

These activities undertaken by TWIST1 during lineage commitment are directed by E-proteins. E-proteins display overlapping expression patterns and developmental functions in embryonic tissues (Murakami *et al*., 2004; Wang and Baker, 2015). They are expressed in the cranial mesenchyme and proliferating neuroepithelium at mid-gestation (E9.5-E11.5) (Murakami *et al*., 2004). *Tcf3* and *Tcf12* expression are restricted to the proliferative ventricular and sub-ventricular zones and are down-regulated during neurogenesis (Uittenbogaard and Chiaramello, 2002; Pfurr *et al*., 2017). *Tcf4* expression persists in the adult brain in regions that overlap with regions expressing genes encoding neuronal bHLH factors such as *Atoh1, Ascl1* and *Neurog2*, and initiates neuronal specification (Forrest *et al*., 2014). Consistent with these findings, tethered TWIST1-TCF4 was shown to modulate the activity of neural-ectoderm genes, whereas tethered TWIST1-TCF3 most strongly activated the mesoderm genes. TWIST1-E-protein heterodimers activated the neural crest cell differentiation and TGFβ signalling pathway, which in turn up-regulated ID proteins that could compete for E-protein binding, providing a potential negative feedback mechanism to modulate the activity of the heterodimers (Connerney *et al*., 2006).

In conclusion, our study has uncovered the predominant dimerization partners of TWIST1 and their role in regulating lineage differentiation during early development. TWIST1 interacts with TCF proteins to promote the differentiation of neural crest and mesoderm progenitors, and to repress endoderm lineage differentiation with the help of TWIST1 homodimer. The TWIST1-TCF dimers enable the mesenchyme precursors to acquire myogenic, skeletogenic and neurogenic fates. In the context of embryogenesis, we showed that the mutations of *TWIST1* in congenital diseases are likely to alter the balance of dimer compositions, which may be responsible for the mesenchymal tissue defects observed in adults.

## Materials and Methods

### Cell lines and cell culture conditions

Cell lines and their treatments are summarized in Table S6. The Madin-Darby Canine Kidney (MDCK) cell line over expressing human *TWIST1* was received from Xue and colleagues (Xue *et al*., 2012). Cells were maintained in high glucose Dulbecco’s modified Eagle’s medium (DMEM, Gibco), 10% Fetal calf serum (FCS) and 10 mM β-mercaptoethanol at 37°C and 5% CO2. *TWIST1* expression was under constant selection using 4 μg/mL of puromycin (Xue *et al*., 2012). C3H10T1/2 cell line was received from ATCC. Cells were maintained in high glucose DMEM with 10% FCS (Life Tech, Australia) and 10 mM β-mercaptoethanol at 37°C in 5% CO2.

Mouse ESCs (A2loxCre) was a gift from Kyba Lab (Lillehei Heart Institute, Minnesota, USA), originally described in Mazzoni *et al*., (2011). The ESCs were cultured in complete mESC medium (high glucose DMEM (Gibco), 12.5 % (v/v) heat-inactivated FBS (Fisher Biotec Australia), 10 mM β-mercaptoethanol, 0.09 % (v/v), 1X non-essential amino acids (100X Thermo Fisher Scientific), 1 % (v/v) nucleosides (100X, Merck) and 10 mil U/mL ESGRO^®^ mouse leukaemia inhibitory factor (Merck)).

All cell lines were tested free of mycoplasma.

### Generation of *Twist1* mutant ESCs by CRISPR-Cas9

ESC editing and selection by GFP was performed as described in detail in (Sibbritt *et al*., 2019). Briefly, 3µg of pSpCas9(BB)-2A-GFP (addgene plasmid #48138, a gift from Feng Zhang) expressing the gRNA of interest were electroporated into 1×10^6 A2loxCre ESCs using the Neon^®^ Transfection System (Thermo Fisher Scientific). Colonies expressing GFP were picked 4 days after electroporation.

For genotyping, region surrounding the mutation site (+/- 200 bp) was amplified from cell lysate, purified and cloned for sequencing. Clones with mono- or bi-allelic frameshift mutations were expanded and used for subsequent experiments.

### Generation of dimer inducible ESC line

The inducible TWIST1-dimer ESCs lines were generated using the inducible cassette exchange method described previously (Mazzoni *et al*., 2011; Iacovino *et al*., 2014). The protein coding sequences were cloned from mouse embryos cDNAs into the p2lox plasmid downstream of the Flag-tag coding sequence (Mazzoni *et al*., 2011; Iacovino *et al*., 2014). Plasmids were transfected into A2loxCre treated with 1 μg/mL doxycycline for 24 hrs. Selection was performed in 300⍰μg/mL of G418 (Gibco) antibiotic for one week. Colonies were then picked and tested for constructs expression following doxycycline treatment.

### Embryoid body formation

Embryoid bodies (EB) were maintained in EB-medium consisting of high glucose DMEM supplemented with 15% heat-inactivated FCS, 1% Non-essential amino-acid supplement and 10 mM β-mercaptoethanol. Single ES cells were seeded onto aggrewells in a 24 well plate (Stem Cell Technologies, Cat. No. 34411) at 1.5 × 10^6 cells per well in EB-medium. Cell aggregates were transferred to non-tissue culture treated 10 cm plates and EB-medium 24 h later and placed onto an orbital-shaker under slight rotation at 37 °C, 5% CO2. Drug treatment conditions was optimised as shown in Supplemental Fig. S2. Doxycycline was supplied at 50 ng/mL for T3 and T12, 100 ng/mL for TT, and 1000 ng/mL for other cell lines after 2 days in differentiation media. Cells were collected 16 hours later as expression of known *TWIST1* targets peaked 12–16 hrs after induction (Supplemental Fig. S2B).

### *In situ* hybridisation

The use of C57BL/6-Arc E 9.5 mouse embryos was approved by the Children’s Medical Research Institute/Children’s Hospital at Westmead Animal Ethics Committee. Embryos collection and in situ hybridisation were performed as previously described (Bildsoe *et al*., 2016).

### Scratch Assay

TWIST1*-CC155 (1 μg) and TCF12-CrN173 (1 μg) were co-transfected into C3H10T1/2 cells seeded in wells of 6-well plate. 24 h after transfection and when cells had reached confluency, a p200 pipet tip was used to make a controlled scratch on the cell monolayer. Cells were then washed once with cell culture medium to remove floating cells detached by the scratch. Plates were then installed into the chamber (37°C, 5% CO2) of a Cell Observer Widefield microscope (ZEISS international) and bright-field images of the region scratched were taken every 15 min over a 8 h period. ImageJ was used to process the imaging data. Briefly, images were enhanced by subtracting the background and binary images were created to assist quantification of the gap closure. Total migration from the start of imaging until the time point when the first cell line closed the gap was then calculated. One-way ANOVA followed by wo-tailed t-test was used to determine the significance of the difference observed between the different conditions.

### Protein Affinity Purification and Mass spectrometry

2×10^7 *MDCK/hTWIST1* over-expressing cells were collected for each experiment. Cell pellets were thawed in 500 μl hypotonic lysis buffer (HEPES 20 mM, MgCl2 1 mM, Glycerol 10%, Triton 0.5%, DTT 1 mM, 1X Complete protease inhibitor [Roche], 1000 U/ml Benzo nuclease) and incubated at room temperature for 15 min. Equal volume of hypertonic lysis buffer (HEPES 20 mM, NaCl2 500 mM, MgCl2 1 mM, Glycerol 10%, DTT 1 mM, 1X Complete protease inhibitor) was applied to the lysate. Cells were further broken down by passage through gauge-25 needle 10 times and then rotated at 4°C for 30 min. After centrifugation at 14,000 x g, 15 min, 450 μL lysates were incubated with 5 µg of an α-TWIST1 antibody or mouse IgG at 4°C overnight with rotation. Meanwhile, Dynabead^®^ protein G slurry (Invitrogen™, cat. 10003D) was prepared by washing and blocking for 1 h at room temperature in 1% BSA 0.1% triton dissolved in PBS 1X before use. 50 μL of Dynabead was then added to each sample and rotated at room temperature for 30 min. Beads were then washed with 500 μL ice-cold wash buffer (1:1 mixture of the lysis buffer, without nuclease) 6 times and transferred to new 1.5 ml tube before the last wash. Beads were then washed quickly with 1 mL cold Tris·HCl pH 7.4 50 mM and then 500 μl TEAB (75 mM). Lastly, beads were collected by centrifugation for 5 min at 2,000 × *g*, before being processed for Mass-spectrometry analysis.

Tryptic digestion of the immunoprecipitated proteins was performed by adding trypsin 1:20 (µg) directly to the washed beads in 50 mM TEAB buffer, vortex and incubation at 37 °C overnight. A second digestion was performed by adding trypsin 1:40 the next day and incubation for 4 h. The tubes were then placed on a magnet and the supernatant was collected and directly transferred into TFA to obtain a final concentration of 0.5%.

Proteolytic peptides were desalted using Oligo R3 reversed phase resin (Thermo Fisher Scientific) in custom-made stage tips (Rappsilber *et al*., 2007). Mass spectrometry (MS) was performed using an LTQ Velos-Orbitrap MS (Thermo Fisher Scientific) coupled to an UltiMate RSLCnano-LC system (Thermo Fisher Scientific). Raw MS data files were processed using Proteome Discoverer v.1.3 (Thermo Fisher Scientific). Processed files were searched against the UniProt mammalian database (downloaded Nov 2016) using the Mascot search engine version 2.3.0. Searches were done with tryptic specificity allowing up to two missed cleavages and a tolerance on mass measurement of 10 ppm in MS mode and 0.3 Da for MS/MS ions. Using a reversed decoy database, a false discovery rate (FDR) threshold of 1% was used. The lists of protein groups were filtered for first hits.

### Rank Product analysis for MS data

We generated three sets of data using the non-parametric rank product method (Breitling *et al*., 2004) and based on how many times a protein was reported across independent experiments (Supplemental Table S2). To test for cases where proteins were preferentially detected (here “preferentially” is defined as present in at least 2 samples for Twist, i.e. Set 1, and 2 samples for Control, i.e. Set 2, and not detected in at least 2 samples of their respective contrast), we employed a single class test. If proteins were present in at least two Twist and two control samples, we deemed this set suitable for a two-class comparison, i.e. direct comparison of protein levels (set 3). The overall schema is represented in Fig. 1E and summarised below:

Set 1: present preferentially in IPs (n ≥ 2)

Set 2: present preferentially in control IPs (n ≥ 2)

Set 3: present in Twist and control cell IPs (n ≥ 2)

P-values and FDR were then determined using the rank product method implemented in the RankProd R package (version 2.34) using 1000 permutations (Hong *et al*., 2006). We used an FDR threshold of 0.20 in each set. If a protein for Set 3 was present in Sets 1 or 2 it was removed after FDR calculation.

### Protein Immunoprecipitation

For the validation of protein interaction, hTWIST1 expressing MDCK cells were transfected 24 h before immunostaining using Lipofectamine^®^ 3000 (Life Tech) according to manufacturer instructions with one of the following plasmids: *pCMV-gfp-HA, pCMV-Tcf3-HA, pCMV-Tcf4-HA, pCMV-Twist2-HA, pCMV-Ebf1-HA, pCMV-Ebf3-HA, pCMV-Sim2-HA, pCMV-Tal1-HA*.

Cell pellets were thawed in 300 μL hypotonic lysis buffer (HEPES 20 mM, MgCl2 1 mM, Glycerol 10%, Triton 0.5 %, DTT 1 mM, 1X Complete protease inhibitor [Roche], 1000 U/ml Benzo nuclease) and incubated at room temperature for 15 min. Equal volume of hypertonic lysis buffer (HEPES 20 mM, NaCl2 500 mM, MgCl2 1 mM, Glycerol 10%, DTT 1 mM, 1X Complete protease inhibitor) was applied to the lysate. Cells were further broken down by passage through a gauge-25 needle 10 times and rotated at 4°C for 30 min before to be centrifuged at 14,000 x g for 15 min. The supernatant was incubated with an α-TWIST1 or an α-FLAG antibody (2 ug/ml) at 4°C for 2 h. 1/10 volume of Protein-G agarose beads (Roche) was then added and the sample rotated for 30 min at room temperature. Beads were finally washed with ice-cold wash buffer (1:1 mixture of the two lysis buffers) 6 times and transferred to a new 1.5 ml tube before. Samples were eluted in 2x LDS loading buffer (Life Technologies) at 70°C for 10 min and half of the eluate was loaded on SDS-PAGE in parallel of the “input” control for western blot analysis.

### Western Blotting

Proteins were extracted using RIPA lysis buffer (1X PBS, 1.5% Triton X-100, 1% IGEPAL, 0.5% Sodium Deoxycholate, 0.1% SDS, 1 mM DTT, 1X Complete protease inhibitor [Roche]) for 30 minutes at 4°C while rotating. Lysed samples were then centrifuged at 15000 x g and the supernatant was collected. Protein concentration was measured using the Direct Detect spectrometer (Millipore). 20 µg of protein was denatured at 70°C for 10 min in 1X LDS loading buffer. Protein electrophoresis and transfer was performed using the NuPage system (Life Technologies, Cat. NP0322BOX), following manufacturer instructions.

Primary antibodies used were mouse monoclonal anti-TWIST1 (1:1000, abcam, Cat. #ab50887), mouse anti-α-tubulin (1:1000, Sigma, Cat. #T6199), rabbit anti-HA (1:1000, abcam, Cat. #ab9110), mouse anti-FLAG M2 (1:1000, Sigma, Cat. #F1804), mouse anti-TCF4 (1:500, Santa Cruz, Cat. #sc-166699), rabbit anti-TCF12 (1:400, Santa Cruz, Cat. # SC-357 1) and mouse anti-GFP (1:1000, Thermo Fisher Scientific, Cat. # A-11120). Secondary antibodies used were HRP-conjugated donkey anti-Rabbit IgG (1:8000, Jackson Immunoresearch, Cat. #711-035-152) and HRP-conjugated donkey anti-Mouse IgG (1:8000, Jackson Immunoresearch, Cat. #711-035-150).

### RNA-seq and data analysis

The raw RNA-seq sequence data were deposited into the NCBI GEO database and can be accessed with the accession number GSE130252. Token for data access: elajseaoxnmbtwv.

RNA library prep (Truseq mRNA kit) and sequencing service (Hiseq2500, 100bpPE) were performed by Macrogen (Seoul, Korea). Raw Fastq sequence files (20 Million reads per sample) were imported into Galaxy (https://galaxy-mel.genome.edu.au/) and analysed using a standard pipeline outlined in Fig. 3.8 (Anders *et al*., 2013). FastQC package was used to assess data quality and adapter sequences and low-quality reads were trimmed from the original reads (Slidingwindow method in Trimmomatic package). A minimum of 50 bp read length filter was applied. Alignment was performed using Tophat2 package using the built-in mm10 genome. The resulting bam file was sorted and assessed visually before reads were counted using the htseq-count package and Genecode mouse basic genome assembly (GRCm38, vM10).

Read count values were imported into the R software for DEG analysis (Afgan *et al*., 2016)). Genes that were not expressed at a biologically meaningful level under any conditions were discarded. A standard count per million (CPM) reads value threshold of 1 (or log-CPM value of 0) was used for considering a gene to be expressed. To be kept for downstream analysis, a gene had to be expressed in at least three samples across the entire experiment. After filtering using this criteria, the number of genes was reduced to 14,165, approximately half the original number. One replicate of T12 sample was removed from further analysis due to low *Tcf12* induction level, and abnormal sample distance from other replicates in correlation map. For EdgeR and limma-voom analysis, the data were normalised by library size before further processing.

Pre-processed data were analysed with R/Bioconductor packageDESeq2, (Love *et al*., 2014). The common adjusted P-value (adjp) threshold of 0.05 and Log2FC of 2 was used. Two types of gene-set analysis were applied: competitive gene-set analysis using clusterProfiler R package (Yu *et al*., 2012) in Figure 5G, and self-contained gene-set analysis using ROAST package in Table S2 and summarised in Supplemental Fig. S4 (Wu *et al*., 2010). In contrast to commonly used competitive gene-set analysis, the self-contained test takes gene expression into account and examines whether the genes in the set/pathway are differentially expressed as a whole for different dimers. Motif analysis on DEGs was performed using the Homer package (Heinz *et al*., 2010).

### Plasmid construction

Plasmids used in the BiFC experiments were a generous gift from Hu Lab and imported via the Addgene depository. The plasmid backbone and the describing publication are listed in Table S3 (Shyu *et al*., 2006). *Twist1* and *Tcf12* coding sequences were cloned from cDNAs generated from E9.5 mouse head embryos and inserted into the relevant expression plasmids. Mouse homologous of S123A, R118H, T121E;S123E, T148A, A152P and I156N variants were generated by site-directed mutagenesis as described previously (Liu and Naismith, 2008) using primers listed in Supplemental Table S5.

### Bi-Fluorescent Complementation (BiFC) Experiments

BiFC experiments and quantitative analysis were performed as described previously (Hu and Kerppola, 2003). The day before transfection, C3H10T1/2 cells were seeded on 0.1% gelatine coated coverslips at a density of 1.66×10^6 cells per 9.6 cm^2^ (well of 6-well plate). 24 h later, transient transfection was performed using Lipofectamine 3000 (Life Technologies) according to manufacturer’s instructions. Equimolar quantities (1 μg each) of plasmids encoding TWIST1-VN173, TCF12-CrN173 and wildtype or mutant TWIST1-CC155 (Fig. S1A) were co-transfected.

Cells were imaged 16-24 hours following transfection. Before imaging, cells were brought to room temperature, washed twice with PBS 1X and fixed in 4% paraformaldehyde (dissolved in PBS 1X) for 10 min. Coverslips were then washed three times with PBT (1% Tween 20 in PBS 1X), once with 1% Trition X-100 in PBS 1X for 5 min and three times with PBT. This was followed by incubation with 4’,6-diamidino-2-phenylindole (DAPI; Sigma-Aldrich; 1:10000) for 10 min at room temperature. Coverslips were then washed three times with PBT for 5 min in the dark and mounted using Fluoromount-G (Invitrogen).

All immunofluorescence slides were imaged using Zeiss Axio Imager A1 (Carl Zeiss, Australia). TWIST1*-CC155/TCF12-CrN173 heterodimers were visualized using a CFP filter set. A GFP filter set was used for detection of TWIST1*-CC155/TWIST1-VN173 homodimers. DAPI was imaged at 405 nm. For BiFC quantification, regions of the slides to be imaged were randomly chosen and pictured at 20 X magnification. At least 1000 cells for each treatment (based on DAPI counting) were sampled across three biological replicates. Fluorescence intensity for the nucleus (DAPI stained) was quantified using ImageJ software and analysed in R (scripts are available upon request).

After filtering for background fluorescence, using the same intensity threshold for all treatment groups, the ratio between CFP/GFP was taken for each cell. Mean value of this ratio was used to compare the CC155/CrN173 and CC155/VN173 dimer formation efficiencies between wildtype or mutant TWIST1. Non-parametric Mann-Whitney U Test was used for statistical analysis. TWIST1* represents mouse homologous of wildtype, S123A, R118H, T121E; S123E, T148A, A152P or I156N variants.

### Web Resources

OMIM: http://www.omim.org/

Galaxy Australia: https://usegalaxy.org.au/

The Human Gene Mutation Database (HGMD^®^): http://www.hgmd.cf.ac.uk/

AQUARIA: http://aquaria.ws

## Acknowledgements

Imaging analysis was performed at the ACRF Telomere Analysis Centre and proteomics analysis was performed at the Biomedical Proteomics Facility, both supported by the Australian Cancer Research Foundation. Our work was supported by the Australian Research Council (DP 1094008, DP 160100933) and Mr James Fairfax (Bridgestar Pty Ltd). XF was supported by the University of Sydney International Postgraduate Research Scholarship, the Australian Postgraduate Award and CMRI Scholarship. PPLT is a NHMRC Senior Principal Research Fellow (Grant ID 1110751).

## Author contribution

P.P.L.T., N.F. and X.F. designed the project; X.F., M.D. and J.S. conducted the experiments; M.G. and D.L. provided technical assistance with proteomics and transcriptome analysis; X.F., A.W. and P.O. performed the bioinformatics analysis; X.F., A.W., N.F. and P.P.L.T. wrote the manuscript.

## Declaration of interest

The authors declare no competing interests

## Notes

#### Summary of Updates

Original Figure 1& 2 are combined; Validation data added to Figure 5; Example imaging data added to Figure 2; Comprehensive analysis added to highlight different functions of dimers on Figure 4; Clarity and readability of the text has been improved;

