## Supplemental Figures S1-S6 for "TWIST1 homodimers and heterodimers orchestrate lineage-specific differentiation"

### Supplementary Tables

Table_S1 bHLH factors expressed in E 9.5 mouse head

Table_S2 Analysis of mass-spectrometry data

Table_S3 Dimer DEG and Gene-set analysis

Table_S4 Gene-set analysis T12 vs TT cell lines

Table_S5 Primers and Plasmids

Table_S6 Summary of cell lines and treatments used in this study

**A**

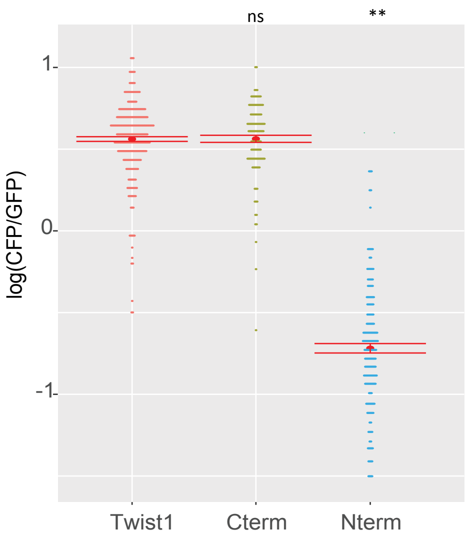


**B**

| Human Variant | | Mouse Variant | |
|---|---|---|---|
| Protein | Gene | Protein | Gene |
| R118H | *353G>A* | R122H | *365G>A* |
| S123A | *367T>G* | S127A | *379T>G* |
| T121E;S123E | *361A>G* | T125E;S127E | *373A>G* |
| | *362C>A* | | *374C>A* |
| | *367T>G* | | *379T>G* |
| | *368C>A* | | *380C>A* |
| T148A | *442A>G* | T152A | *454A>G* |
| A152P | *454G>C* | A156P | *466G>C* |
| I156N | *467T>A* | I160N | *479T>A* |

**Figure S1.** Additional information for BiFC experiments. **A.** Heterodimer/homodimer CFP/GFP ratio quantified for individual cells (each dot corresponding to one cell) for full-length TWIST1, the C-terminal region (Cterm, contains bHLH domain) and the N-terminal region (Nterm) of TWIST1. **B.** Human-mouse correspondence for variants used in this study.

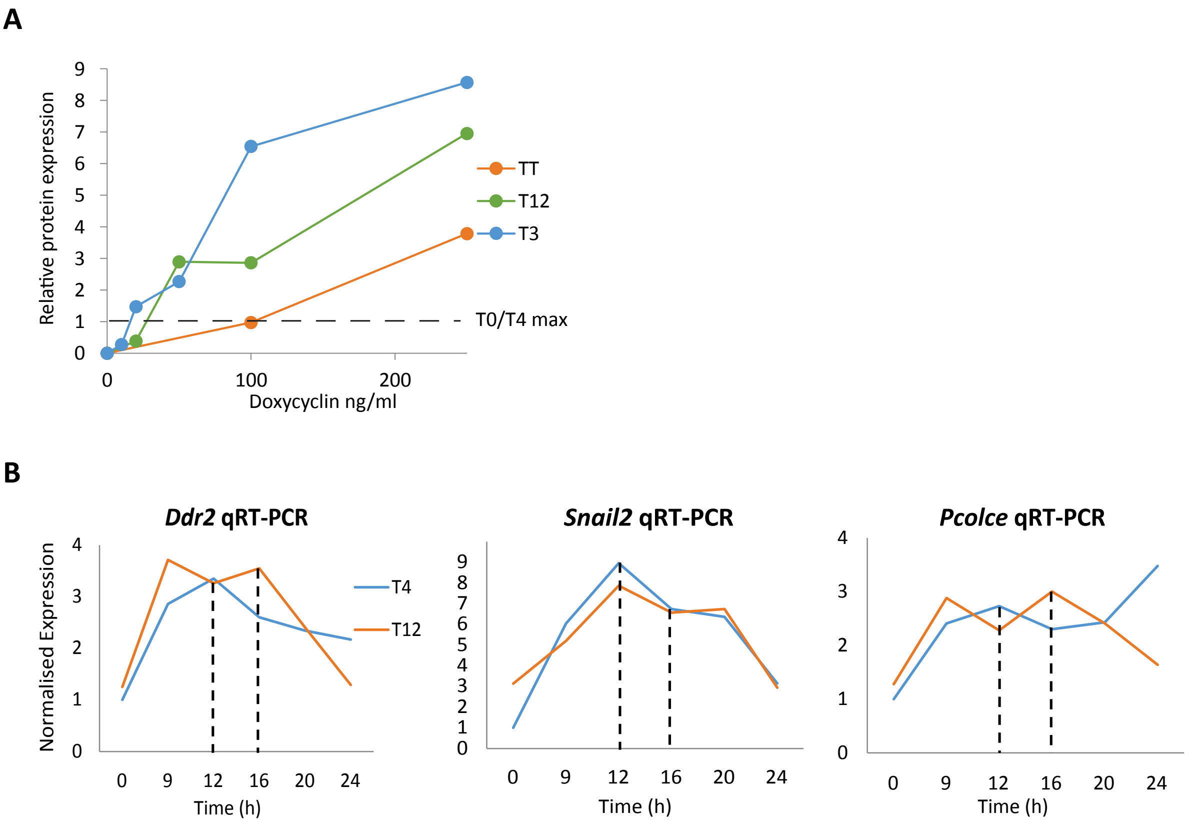


**Figure S2**. Optimisation of doxycycline dosage to induce comparable levels of expression of the constructs between the different engineered ESCs cell lines. **A.** Preliminary analysis (not shown) determined that level of expression (Y-axis) of T4 and T0 constructs were maximal and similar for 1000 ng/ml of doxycycline (dotted line on the graph). Different concentrations of doxycycline (X-axis) were tested for cell lines expressing TT, T12 and T3 constructs to identify which concentrations are required to achive comparable levels of expression between all the cell lines. Protein levels were quantified by western blot using an α-TWIST1 antibody and normalised against β-Tubulin expression, after 16 h of exposure to doxycycline. **B.** RT-qPCR analysis of the expression TWIST1 direct target genes Ddr2, Snail2 and Pcolce relative to Actb expression (Y-axis) at different time points after doxycycline treatment (X-axis) in cell lines expressing T4 and T12 constructs.

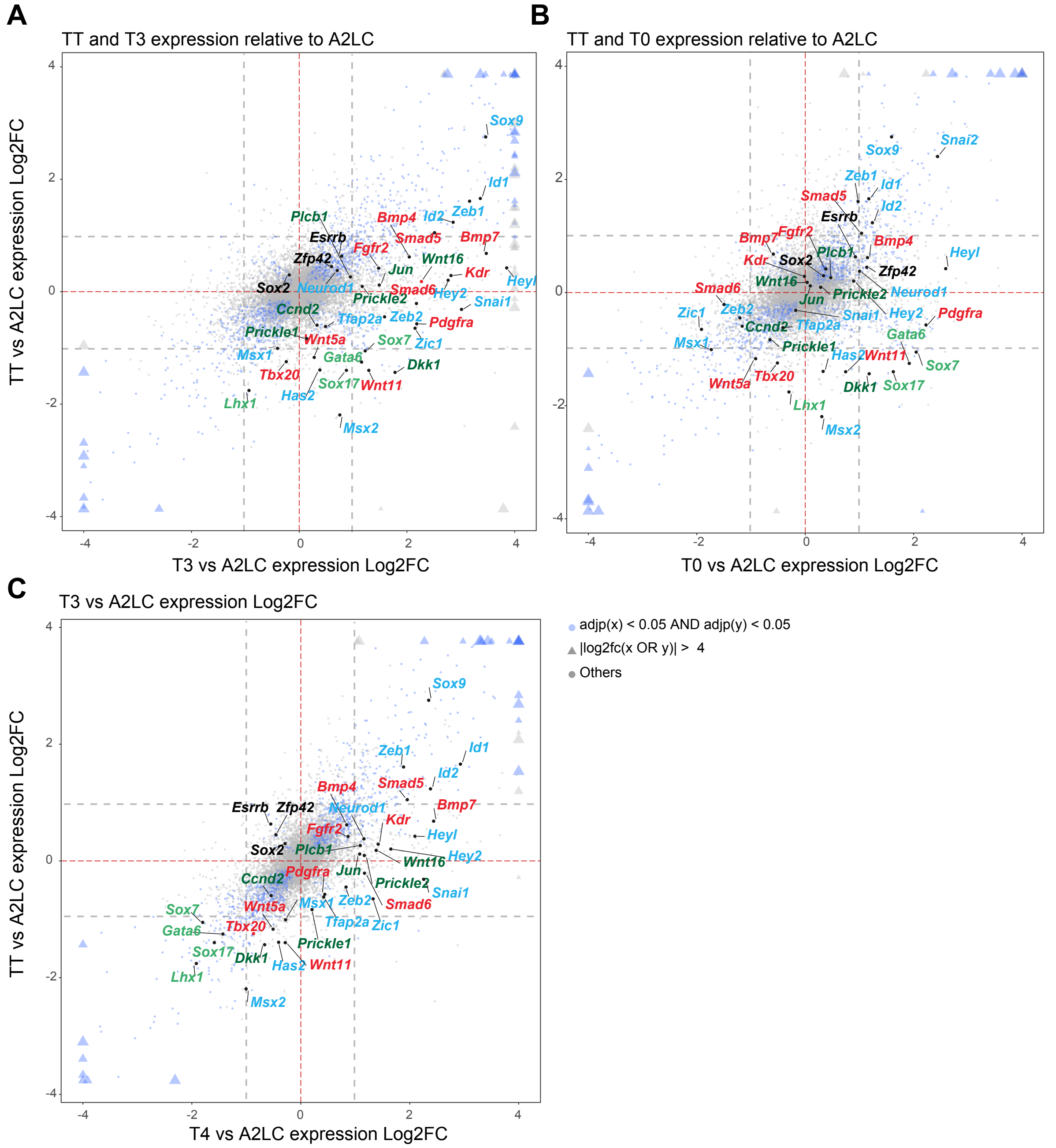


**Figure S3.** Dimer specificity comparison. **A-C.** Four-way plot visualizations with DESeq2 data. Log2FCs comparisons between TT versus A2LC, and T3 or T4 versus A2LC cell lines are made. Blue nodes indicate statistically significant genes for both cell lines. Triangular shapes indicate values which exceed the viewing are for the graph. Vertical and horizontal dashed lines indicate |Log2FC|= 1. Size change reflects the magnitude of Log2FC values. Vertical and horizontal dashed lines indicate |Log2FC|= 1. Key markers and pathway genes are indicated: Black, pluripotency; Green, endoderm; Red, mesoderm; Blue, ectoderm & EMT; Dark green, WNT signalling pathway.

**A**

| T0 | TT | T3 | T4 | T12 |
|---|---|---|---|---|
| Cardiac Atrium Development | Cardiac Atrium Development | Artery Development | Dendrite Development | Cardiac Septum Development |
| Olfactory Lobe Development | Endoderm Development | Coronary Vasculature Development | Regulation Of Erythrocyte Differentiation | Dendrite Development |
| Endocrine Pancreas Development | Placenta Blood Vessel Development | Myeloid Leukocyte Differentiation | Face Development | Cerebral Cortex Development |
| Pallium Development | Mesoderm Development | Cardiac Septum Development | Neuron Projection Development | Positive Regulation Of Dendrite Development |
| Hippocampus Development | Embryonic Skeletal System Development | Regulation Of Skeletal Muscle Tissue Development | Regulation Of Epidermal Cell Differentiation | Cardiac Chamber Development |

**B**

| Gene ontology | Ngenes | Propdown | Propup | Direction | Pvalue | FDR |
|---|---|---|---|---|---|---|
| Neuron Differentiation | 814 | 0.06 | 0.54 | Up | 0.001 | 0.002 |
| Head Development | 667 | 0.05 | 0.55 | Up | 0.003 | 0.006 |
| Vasculature Development | 450 | 0.07 | 0.54 | Up | 0.001 | 0.002 |
| Heart Development | 448 | 0.05 | 0.63 | Up | 0.001 | 0.002 |
| Skeletal System Development | 433 | 0.05 | 0.50 | Up | 0.002 | 0.004 |
| Forebrain Development | 341 | 0.06 | 0.55 | Up | 0.002 | 0.004 |
| Eye Development | 308 | 0.06 | 0.52 | Up | 0.003 | 0.006 |
| Urogenital System Development | 280 | 0.06 | 0.50 | Up | 0.001 | 0.002 |
| Muscle Organ Development | 254 | 0.07 | 0.53 | Up | 0.001 | 0.002 |
| Muscle Tissue Development | 253 | 0.05 | 0.55 | Up | 0.004 | 0.007 |

**C**

| Molecular pathways | Ngenes | Propdown | Propup | Direction | Pvalue | FDR |
|---|---|---|---|---|---|---|
| Reactome Signaling By GPCR | 477 | 0.071 | 0.382 | Up | 0.004 | 0.015 |
| Calcium Signaling Pathway | 165 | 0.042 | 0.442 | Up | 0.002 | 0.013 |
| Chemokine Signaling Pathway | 155 | 0.065 | 0.477 | Up | 0.006 | 0.017 |
| Wnt Signaling Pathway | 141 | 0.043 | 0.645 | Up | 0.006 | 0.017 |
| Reactome Signaling By PDGF | 117 | 0.043 | 0.684 | Up | 0.004 | 0.015 |
| Plasari Tgfb1 Signaling | 100 | 0.030 | 0.740 | Up | 0.005 | 0.016 |
| Signaling Of Activated FGFR | 95 | 0.074 | 0.611 | Up | 0.006 | 0.017 |
| Reactome Signaling By Notch | 93 | 0.054 | 0.656 | Up | 0.005 | 0.016 |
| Kobayashi EGFR Signaling 24hr Up | 85 | 0.059 | 0.482 | Up | 0.004 | 0.015 |
| TGF Beta Signaling Pathway | 83 | 0.060 | 0.614 | Up | 0.001 | 0.007 |

**Figure S4.** Analysis of dimer-specific target genes. **A.** Gene ontology enrichment analysis of genes induced by TWIST1dimer and TWIST1 monomer against the parental A2LoxCre ESCs. Gene sets were filtered by total gene number > 20, p-value < 0.05. The top five significant gene ontologies are indicated. Black, upregulated sets; Red, downregulated sets. **B, C.** Gene ontology and molecular pathways enrichment analysis of genes induced by T12 versus TT. Gene sets were filtered same as A.

**Table S6**. Summary of cell lines and treatments used in this study

<table>
<tr><th colspan="2">Cell lines</th><th>Description</th><th>Treatments</th><th>Experiment</th></tr>
<tr><td colspan="2">MDCK (hTWIST1 O/E)</td><td>MDCK cell stably overexpressing hTWIST1 (Xue et al., 2012)</td><td>Transfection with pCMV-XX-HA constructs for partners</td><td>IP mass-spec; CoIP validation</td></tr>
<tr><td colspan="2">C3H10T1/2</td><td>Mesenchymal stem cell (ATCC)</td><td>Transfection with BiFC constrcts</td><td>BiFC; Scratch Assay</td></tr>
<tr><td rowspan="8">mESC</td><td>A2loxcre</td><td>Mazzoni et al., 2011</td><td rowspan="8">EB formation and Doxycyclin induction</td><td rowspan="8">qPCR analysis or RNA-seq</td></tr>
<tr><td>Twist1-/-</td><td rowspan="2">Derived from A2loxcre by CRISPR editing</td></tr>
<tr><td>Twist1+/-</td></tr>
<tr><td>T0</td><td rowspan="5">Derived from A2loxcre by induced casset exchange; Dox inducible</td></tr>
<tr><td>TT</td></tr>
<tr><td>T3</td></tr>
<tr><td>T4</td></tr>
<tr><td>T12</td></tr>
</table>
